# Efficient Finite-Difference Estimation of Second-Order Parametric Sensitivities for Stochastic Discrete Biochemical Systems

**DOI:** 10.1101/2024.11.04.621989

**Authors:** Fauzia Jabeen, Silvana Ilie

## Abstract

Biochemical reaction systems in a cell exhibit a stochastic behaviour, owing to the unpredictable nature of the molecular interactions. The fluctuations at the molecular level may lead to a different behaviour than that predicted by the deterministic model of the reaction rate equations, when some reacting species have low population numbers. As a result, stochastic models are vital to accurately describe the system dynamics. Sensitivity analysis is an important method for studying the influence of the variations in various parameters on the output of a biochemical model. We propose a finite-difference strategy for approximating second-order parametric sensitivities for stochastic discrete models of biochemically reacting systems. This strategy utilizes adaptive tau-leaping schemes and coupling of the perturbed and nominal processes for an efficient sensitivity estimation. The advantages of the new technique are demonstrated through its application to several biochemical system models with practical significance.

## 1 Introduction

Modelling and simulation of biochemical interactions among molecular species in a living cell are some important topics in Computational Biology. These molecular interactions are inherently random and discrete in nature. Thus, stochastic models are necessary to capture this intrinsic stochasticity, and provide a more accurate representation of cellular processes. Such models are particularly important when some molecules involved in key biochemical reactions are present in low numbers. Stochastic models have proven crucial in studying the complex dynamics of biological processes, ranging from gene regulation to cellular signalling [1, 2]. The Chemical Master Equation is a commonly used discrete stochastic model which accurately represents the dynamics of homogeneous biochemical systems, with some reactants in small molecular amounts [3]. The solution of this model can be approximated numerically using Monte Carlo techniques. Among the most popular Monte Carlo strategies that provide statistically exact trajectories are the stochastic simulation algorithm (SSA) due to Gillespie [4, 5] and the Next Reaction Method developed by Gibson and Bruck [6]. These methods are often expensive on systems with large number of reactive events or with some species in large molecular amounts. Then, approximate Monte Carlo methods such as *τ* -leaping schemes [7, 8] can significantly reduce the computational cost of the simulation while maintaining an excellent accuracy. Other numerical methods for the Chemical Master Equation include [9, 10, 11, 12, 13, 14, 15, 16] (see also references therein).

Sensitivity analysis plays a prominent role in understanding the robustness of biochemical models by quantifying the influence of parameter variations on model outputs [17, 18, 19]. In addition, it is an important tool for parameter estimation [20] and practical identifiability analysis of these models [21, 22]. In the deterministic setting, sensitivity analysis techniques such as partial derivatives or finite differences are commonly employed to compute parameter sensitivities [23].

However, adapting these methods to stochastic discrete models is a difficult task [24]. Several approaches have been proposed for such models. Exact strategies for computing sensitivities using Girsanov measure transform were developed in [25], but may be computationally infeasible for models of moderately large to large systems of biochemical reactions or models which are stiff. Methods offering insights into the properties of probability distributions were proposed in [24, 26, 27]. One of the most popular schemes to estimate parameter sensitivities are the finite-difference strategies based on Monte Carlo simulations of the perturbed and nominal processes [16, 28]. Some existing finite-difference techniques for approximating local sensitivities in stochastic biochemical reaction networks depend on simulations utilizing exact Monte Carlo schemes, including the SSA for the Common Random Number [29], the Random Time Change algorithm employed by the Common Reaction Path [29], or the Next Reaction Method used by Coupled Finite Difference (CFD) [28] schemes. Other finite-difference methods rely on approximate Monte Carlo strategies which are appropriate for systems with various degrees of stiffness such as the Coupled Tau-Leaping [30] and the Coupled Implicit Tau-leaping [31] methods.

These techniques were designed to estimate first-order sensitivities for stochastic biochemical models. While first-order sensitivities show the direct influence of parameter variations on the output of the model, second-order sensitivities capture the curvature or nonlinear effects in the system’s response to parameter perturbations. Moreover, the Hessian (second-order sensitivities) plays a central role in optimization, parameter estimation and model validation. If 𝔼[·] represents the expected value, **f** the function of interest, and **X**(*t, c*) the state of the system at time *t*, depending on the vector of *M* parameters *c*, we are interested in computing the second-order sensitivity, *∂*^2^𝔼 [**f** (**X**(*t, c*))]*/∂c*_*j*_*∂c*_*k*_, where *c*_*j*_ and *c*_*k*_ are some parameters of interest, for *j, k* ∈ {1, …, *M*}. Denote by *e*_*j*_ the vector of dimension *M* with the *j*-th component equal 1 and the others 0. Using a forward finite-difference scheme, this sensitivity may be estimated by 𝔼[**f** (**X**(*t, c* + (*h*_*j*_*e*_*j*_ + *h*_*k*_*e*_*k*_)))] − 𝔼[**f** (**X**(*t, c* + *h*_*j*_*e*_*j*_))] − 𝔼[**f** (**X**(*t, c* + *h*_*k*_*e*_*k*_))] + 𝔼[**f** (**X**(*t, c*))] */*(*h*_*j*_*h*_*k*_), with *h*_*j*_ and *h*_*k*_ being small perturbation of the parameters of interest *c*_*j*_ and *c*_*k*_, respectively. It is worth noting that much less research was dedicated to building finite-difference second-order sensitivity estimators for the Chemical Master Equation. The Coupled Finite-Difference method (CFD-2) for approximating the second-order sensitivities for stochastic biochemical systems was proposed in [33]. This method utilizes the modified Next Reaction Method, an exact Monte Carlo strategy, to generate the four correlated processes and is working well on problems which are non-stiff.

When biochemical systems evolve on multiple scales in time, their mathematical models exhibit stiffness [32]. For stiff systems, exact simulation strategies become prohibitively costly. Another limitation of existing coupled finite-difference second-order sensitivity estimators is their lack of scalability when applied to large systems with many reacting species and parameters, which may also be stiff. The computational cost of exact simulations grows rapidly as the system size increases. An important difficulty when designing finite-difference second-order sensitivity estimators is that they are more sensitive to noise than those for first-order derivatives. Thus, strong coupling and reliable Monte Carlo simulation methods are required for an accurate approximation of the second-order derivatives. This paper proposes a novel method to estimate second-order parametric sensitivities of stochastic discrete biochemical models, which relies on an approximate Monte Carlo strategy to generate realizations of the correlated processes **X**(*t, c* + (*h*_*j*_*e*_*j*_ + *h*_*k*_*e*_*k*_)), **X**(*t, c* + *h*_*j*_*e*_*j*_), **X**(*t, c* + *h*_*k*_*e*_*k*_), and **X**(*t, c*). This approximate strategy is based on efficient adaptive *τ* - leaping schemes, expanding the techniques developed by Cao et al. [8]. We utilized a coupling of the four processes above, which ensures that the variance of the sensitivity estimator is reduced, leading to an effective sensitivity approximation. This coupling builds on the foundation of the coupled finite-difference framework [33]. Our tau-leaping based method extends the existing approach by offering improved scalability with the system size and increased efficiency for moderately stiff problems. The ability to take larger time steps and reduce the number of simulated reactions enables our method to handle realistic biochemical systems that would be computationally intensive using finite-difference methods relying on exact stochastic simulation strategies. We note that the Coupled Finite-Difference strategy (CFD-2) considers a perturbation of a certain absolute value for both parameters (*h*_*j*_=*h*_*k*_). If there is a significant difference in the parameter values, comparing their relative importance may become challenging. To avoid this problem, our method defines the perturbations in terms of the relative value *h*, such that *h*_*j*_=*hc*_*j*_ and *h*_*k*_=*hc*_*k*_ are small.

The outline of this paper is given below. In Section 2, the background on stochastic discrete models of well-stirred biochemical networks and their simulation strategies is discussed. Section 3 describes various existing parametric sensitivity techniques and introduces a new finite-difference second order sensitivity estimator for stochastic discrete biochemical networks. Section 4 illustrates the benefits of the proposed sensitivity estimator through numerical tests performed on three models arising in applications. Lastly, Section 5 provides our conclusions.

## 2 Background

### 2.1 Stochastic Modelling of Well-mixed Biochemical Networks

This work focuses on developing tools to analyze a stochastic model that describes the evolution of the number of molecules of some biochemically reacting species confined to a constant volume at a constant temperature. More precisely, we assume that a collection of *N* biochemical species *S*_1_, …, *S*_*N*_, of a homogeneous (well-mixed) biochemical system are interacting through *M* chemical reactions, *R*_1_, …, *R*_*M*_. The state of the system is represented by the vector **X**(*t*) = (*X*_1_(*t*), …, *X*_*N*_ (*t*))^*T*^ at time *t*, wherein each *X*_*i*_(*t*) represents a non-negative integer denoting the quantity of molecules of species *S*_*i*_ at the given time *t*, where *i* = 1, …, *N*. The state vector at the initial time *t*_0_ is given, **X**(*t*_0_) = **x**_0_.

Each reaction channel *R* is defined by its stoichiometric vector *ν*_*ℓ*_ and propensity function *a* (·). Its state change vector is denoted by *ν*_*ℓ*_ = (*ν*_1_, *ν*_2_, …, *ν*_*Nℓ*_)^*T*^, where each element *ν*_*iℓ*_ corresponds to the change in the quantity of the *i*-th molecular species after one reaction *R* fires. Suppose that *X*(*t*) = **x** represents the system state at time *t*, then a single reaction *R*_*ℓ*_ taking place over [*t, t* + *dt*) transitions the system to the system state **X**(*t* + *dt*) = **x** + *ν*_*ℓ*_. The propensity of the reaction *R*_*ℓ*_ is such that *a*_*ℓ*_(**x**)*dt* quantifies the probability of the reaction *R* firing within the time interval [*t, t* + *dt*), under the assumption that *X*(*t*) = **x**.

Define by *P* (**x**, *t*|**x**_0_, *t*_0_) the conditional probability of the system state being **X**(*t*) = **x** at time *t*, provided that the initial state was **X**(*t*_0_) = **x**_0_. The evolution of this conditional probability over time is governed by the Chemical Master Equation [3]

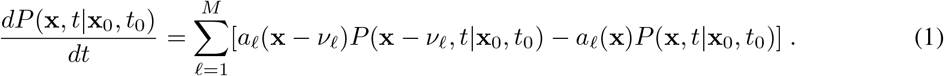

This is a stochastic discrete model of homogeneous biochemical networks. An alternative description of the system state **X**(*t*) which satisfies equation (1) is given by the Random Time Change (RTC) representation (see [34])

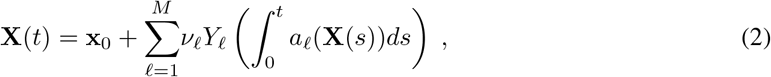

where *Y*_*ℓ*_ with 1≤ *ℓ* ≤ *M* are independent Poisson processes of unit rate and **X**(*t*_0_) = **x**_0_.

For most practical applications, solving equation (1) directly is computationally challenging due to its high dimensionality, in particular for systems with species in large molecular counts and with many reactions. To address this challenge, Monte Carlo strategies can be employed to produce sample paths consistent with the solution of the Chemical Master Equation. An exact Monte Carlo method for generating such realizations of the stochastic process governed by the Chemical Master Equation is the Stochastic Simulation Algorithm (SSA) or Gillespie’s Direct method [4, 5]. The steps of this algorithm are summarized below.

#### Stochastic Simulation Algorithm

1. Initialization: *t* = *t*_0_, *X* = *x*_0_.
2. While *t < T*
3. Calculate *a*_*ℓ*_(*X*) for each *ℓ*, 1 ≤ *ℓ* ≤ *M*, and 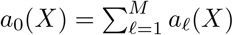.
4. Compute *η*_1_ and *η*_2_, two independent uniform random numbers over [0, 1].
5. Determine the next reaction time 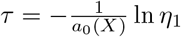.
6. Determine the next reaction index, *j*, with 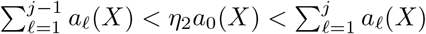.
7. Take *X* = *X* + *ν*_*j*_ and *t* = *t* + *τ*.
8. End while.

Gillespie’s algorithm as well as other exact Monte Carlo methods for the Chemical Master Equation can be computationally intensive for realistic biochemical networks. Simulating individual reaction events sequentially may require substantial computational resources, in particular for systems with some fast reactions.

To achieve a faster computation compared to exact methods, one could utilize approximate Monte Carlo schemes to estimate the solution of the Chemical Master Equation. One such approximate method is the tau-leaping strategy developed by Gillespie [7]. This method may accelerate the simulation significantly for models with multiple scales in time. It does so by advancing the system in time with a predetermined step *τ*, which leaps over many reactions, instead of simulating individual reaction events in a sequential manner. The stepsize is chosen such that the leap condition [7] is satisfied: given that the current system state at time *t* is **X**(*t*) = **x**, the step *τ >* 0 is small enough that each propensity varies insignificantly during [*t, t* + *τ*). With this assumption, the number of reactions *R*_*ℓ*_ taking place within [*t, t* + *τ*) can be approximated by a Poisson random variable with mean and variance equal to *a*_*ℓ*_(**x**)*τ*, namely *P*_*ℓ*_(*a*_*ℓ*_(**x**)*τ*). Therefore the system state may be estimated by

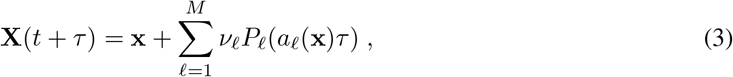

which is known as the (explicit) tau-leaping method [7]. The Poisson random variables {*P*_*ℓ*_}_1≤ *ℓ* ≤*M*_ are independent.

Cao et al. [8] proposed a leap-selection strategy which increases the accuracy and efficiency of the simulation and which is extensively used in applications. To improve the accuracy of the numerical solution, the leap condition is adjusted to require that the relative change in the molecular count of each reactant species, rather than in each propensity, is negligible over the stepsize *τ*. Also, for avoiding negative population numbers, the reacting species are partitioned into critical and non-critical. For this purpose, a control parameter *n*_*c*_ ∈ [2, 20] is introduced. A reaction with a positive propensity is considered critical if *n*_*c*_ occurrences of this reaction drive the molecular amount of one of its reactants negative. Otherwise, the reaction is deemed non-critical. Here *J*_*ncr*_ represents the set of indices of the noncritical reactions, *I*_*r*_ is the set of all reacting species indices and *I*_*ncr*_ is the set of indices of the reacting species participating in non-critical reactions. For a time *t*, a state **X**(*t*) = **x** and a reaction *R*_*ℓ*_ with *a*_*ℓ*_(**x**) *>* 0, the minimum number of firings of that reaction needed to consume one of its reacting species is

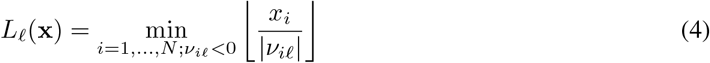

where ⌊·⌋ denotes the greatest integer less than the given value. With this notation, a reaction *R* with a positive propensity is classified as *critical* if *L*_*ℓ*_(**x**) *< n*_*c*_. For any index *i* ∈ *I*_*r*_, consider *r*_*i*_ to be the highest order of a reaction in which the species *S*_*i*_ is a reactant. Then, the functions *g*_*i*_ = *g*(**x**) are defined as follows [8]:

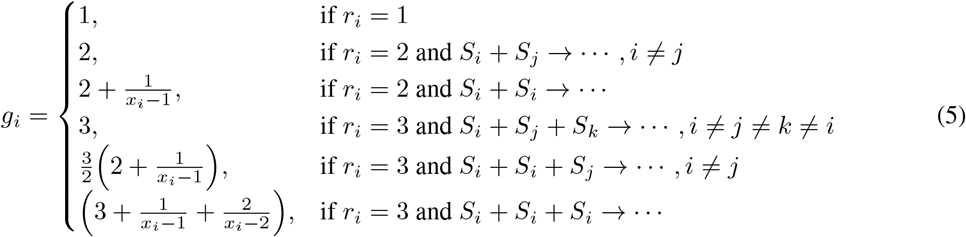

A second control parameter is the tolerance *ε*, which satisfies 0 *< ε <* 1. The modified leap condition requires that the relative change in the molecular counts of each reactant species is (in some sense) below the tolerance *ε*. More precisely, if at time *t*, the state is **X**(*t*) = **x**, then *τ* should be small enough such that

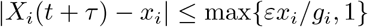

for any *i* ∈ *I*_*r*_. According to [8], the leap size *τ* is computed as

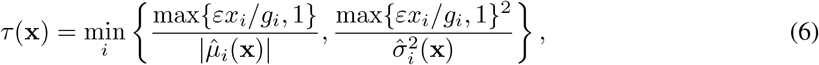

with the auxiliary equations for each species *S*_*i*_ being

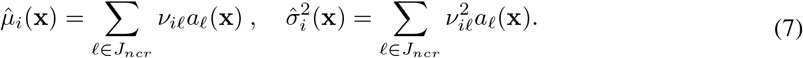

Our strategy for estimating second order sensitivities relies on adaptive tau-leaping methods which utilize *τ* -selection schemes developed in Cao et al. [8].

## 3 Second-Order Parametric Sensitivity

Sensitivity analysis is a fundamental tool in the modelling and analysis of biochemical systems, measuring the impact of perturbations in parameter values on the behaviour of the system. This analysis aims to identify essential parameters that significantly influence the system dynamics, assess the robustness of the model to parameter variations, and provide guidance for experimental design and parameter estimation. Local sensitivity analysis examines the influence of small changes in input parameters on the model outputs. In the case of biochemical systems, these parameters include initial molecular amounts of various species or kinetic parameters. A large local sensitivity in a parameter indicates that the model is sensitive with respect to that parameter, and thus an accurate estimation of its value is necessary.

For stochastic models, the first-order local parametric sensitivity can be computed by *∂*𝔼[**f** (**X**(*t, c*))]*/∂c*_*j*_ where 𝔼[·] symbolizes the expected value, **f** denotes a smooth function of interest, and **X**(*t, c*) is the state of the system at time *t*, corresponding to the vector of parameters *c*, and *c*_*j*_ is the parameter under investigation. Finite-difference techniques are among the most commonly ulitized strategies for estimating these sensitivities. Several finite-difference schemes for approximating first-order sensitivities were proposed in the literature (see, for example, [28, 29, 30, 31]).

In the case of stochastic models, designing effective methods for estimating the second-order parametric sensitivities *∂*^2^𝔼[**f** (**X**(*t, c*))]*/∂c*_*j*_*∂c*_*k*_, where *c*_*j*_ and *c*_*k*_, with *j, k* ∈ {1, *…, M*}, represent two model parameters, is a difficult problem. A forward finite-difference estimation of the second-order sensitivity with respect to these parameters is

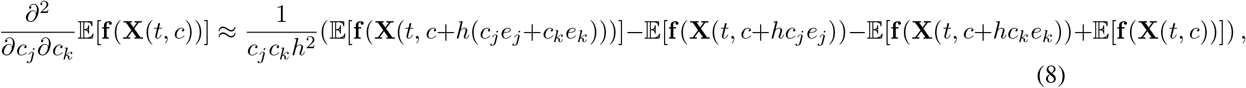

where *h* is a small relative perturbation. The goal is to estimate the expected value of the random variable [**f** (**X**(*t, c* + *h*(*c*_*j*_*e*_*j*_ + *c*_*k*_*e*_*k*_))) − **f** (**X**(*t, c* + *hc*_*j*_*e*_*j*_)) −**f** (**X**(*t, c* + *hc*_*k*_*e*_*k*_)) + **f** (**X**(*t, c*))]*/*(*c*_*j*_*c*_*k*_*h*^2^). Monte Carlo methods may be utilized to generate realizations of this random variable. Then, an estimator of the second-order sensitivity with respect to *c*_*j*_ and *c*_*k*_ is the sample mean of a set of realizations

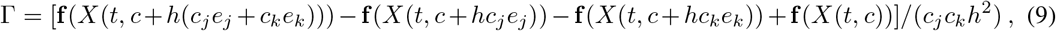

where we denoted by *X*(*t, c*) a sample path of the stochastic process **X**(*t, c*).

When the four processes **X**(*t, c* + *h*(*c*_*j*_*e*_*j*_ + *c*_*k*_*e*_*k*_)), **X**(*t, c* + *hc*_*j*_*e*_*j*_), **X**(*t, c* + *hc*_*k*_*e*_*k*_) and **X**(*t, c*) are correlated, the variance of the sensitivity estimator is reduced. The challenge consists in designing variance reduction schemes which result in a more effective sensitivity approximation. Such a finite-difference strategy based on correlating the four processes was proposed in [33]. For this strategy, an exact Monte Carlo scheme, the Next Reaction Method or Gillespie’s algorithm, is utilized to generate realizations of these processes. This technique estimates second-order sensitivities by ensuring the four paths share reactions, thereby producing low-variance estimates. We shall compare our proposed strategy which shares groups of reactions to this scheme, which is the state-of-the-art method for computing second-order sensitivities for the Chemical Master Equation model.

The four processes, **X**(*t, c* + *h*(*c*_*j*_*e*_*j*_ + *c*_*k*_*e*_*k*_)), **X**(*t, c* + *hc*_*j*_*e*_*j*_), **X**(*t, c* + *hc*_*k*_*e*_*k*_), **X**(*t, c*) have a unified representation:

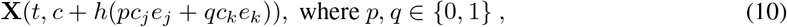

and they share the same initial condition, **X**(0, *c* + *h*(*pc*_*j*_*e*_*j*_ + *qc*_*k*_*e*_*k*_)) = **x**_0_.

Denote the propensity of the reaction *R*_*ℓ*_, 1 ≤ *ℓ* ≤ *M*, of each process by:

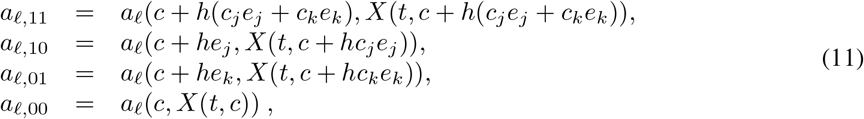

where the notation ignores the dependence on the time *t* and the parameters, for simplicity. Also, consider the set

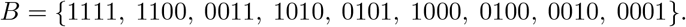

In order to couple the four processes above, which produces a low variance sensitivity estimator, we define the following terms, *α*_*ℓ,b*_, for 1 ≤ *ℓ* ≤ *M* and *b* ∈ *B*, in accordance with [33]:

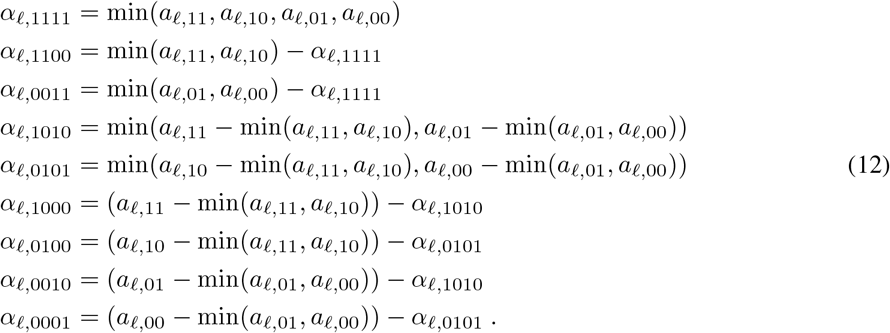

The coupled finite-difference strategy for estimating second-order sensitivities, CFD-2 [33], relies on a strong correlation of the four processes (10). This correlation, which utilizes the RTC representation (2), is provided below:

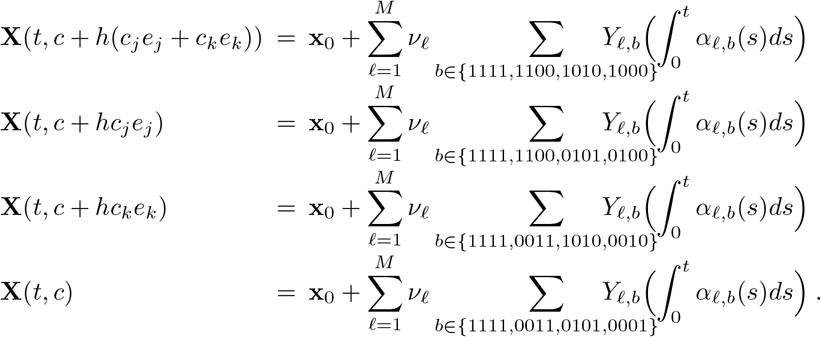

Remark that every counting process *Y*_*ℓ*_ in the RTC representation is split into subprocesses shared among the four continuous time Markov chains (10). Moreover, each of the correlated processes above has the same distribution as that of their uncoupled counterparts. The CFD-2 algorithm utilizes the Stochastic Simulation Algorithm [4, 5] or the Next Reaction Method [28].

### 3.1 Coupled *τ* -Leaping Method for Estimating Second Order Sensitivities

We propose below a strategy which relies on the novel combination of tau-leaping with a finite-difference sensitivity estimation approach that is designed for efficient and accurate second-order parametric sensitivity analysis for stochastic discrete models of homogeneous biochemical networks. We show that second-order sensitivities can be estimated accurately and efficiently by utilizing a variable time-stepping strategy for tau-leaping. This is a valuable and non-trivial extension of an existing finite-difference method, which allows sensitivity analysis to scale efficiently to larger, more complex systems. Also, this technique is effective and reliable for models of biochemical reactions that operate on multiple time scales (stiff problems). Stiff systems are commonly found in practical applications, as biochemical processes typically entail both rapid and slow reactions. For such cases, exact stochastic simulation schemes are computationally intensive, since they advance the system one reaction at a time.

Unlike the coupled finite-difference scheme CFD-2 [33], which utilizes exact methods to generate the coupled quadruple Monte-Carlo trajectories, our approach employs variable *τ* -leaping techniques to compute the four correlated paths. The proposed coupled *τ* -leaping (CTL-2) strategy is based on sharing Poisson random variables among the nominal and perturbed trajectories. Moreover, the impact of the shared terms is anticipated to be substantial, resulting in a strong coupling and thus a notable decrease in the variance of the sensitivity estimation.

In the coupled tau-leaping algorithm for approximating second order sensitivities, the nominal and the three perturbed sample paths are correlated according to

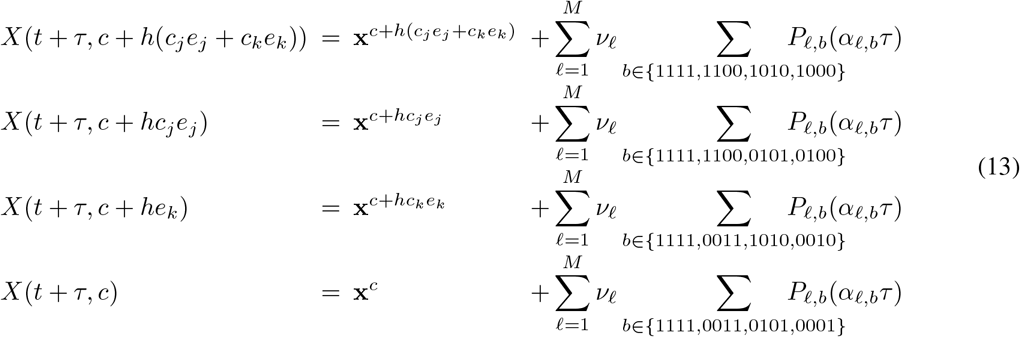

where 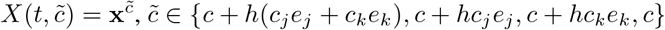. Here *P*_*ℓ,b*_(*α*_*ℓ,b*_*τ*), 1 ≤ *ℓ* ≤ *M* and *b* ∈ *B*, are independent Poisson random variables of mean and variance *α*_*ℓ,b*_*τ*.

To reduce the computational time and avoid negative population numbers in the simulation, we shall extend the variable time-stepping strategy introduced by Cao et al. [8] to generate the four correlated trajectories with the explicit tau-leaping strategy (13). First, a potential stepsize is calculated separately for both critical and non-critical reactions on each of the four trajectories. Then, the smallest of these stepsizes is selected as the next *τ* for the tau-leaping strategy on each path. Note that for the case of double derivatives with respect to one parameter (i.e., *j* = *k*), the paths *X*(*c* + *hc*_*j*_*e*_*j*_) and *X*(*c* + *hc*_*k*_*e*_*k*_) are still generated separately. The steps of the algorithm are given below.

#### Coupled *τ* -Leaping Algorithm

1. **Set model parameters**: tolerance *ϵ*, critical threshold *n*_*c*_, simulation final time *T*, and relative perturbation *h*.
2. **Initialize trajectories** at *t* = 0: *X*(*c* + *h*(*pc*_*j*_*e*_*j*_ + *qc*_*k*_*e*_*k*_)) = **x**_0_, for all *p, q* ∈ *{*0, 1*}*.
3. **While** *t < T* **do** steps (a) – (h):
  a. **Calculate the propensities** from (11): for *ℓ* = 1, …, *M* and *p, q* ∈ *{*0, 1*}*,

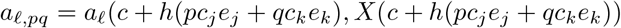
  b. **Find critical reactions** for each of the four trajectories: for each reaction *R*_*ℓ*_ and *p, q* ∈ {0, 1}, if *a*_*ℓ,pq*_ *>* 0, then

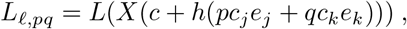

with *L* defined by (4). Put *J*_*ncr*_ = *{ℓ*| *L*_*ℓ,pq*_ ≥ *n*_*c*_ for all *p, q* ∈ *{*0, 1*}}*.
  c. **Find candidate step sizes**, *τ*_*ncr,pq*_, **for the noncritical reactions**; for all *p, q* ∈ *{*0, 1*}*:
    i. If *J*_*ncr*_ = ∅, then *τ*_*ncr,pq*_ = ∞.
    ii. If *J*_*ncr*_ ≠ ∅, then find set of indices of reactant species of non-critical reactions *I*_*ncr*_.
    iii. For each *i* ∈ *I*_*ncr*_ and each trajectory, find the highest order of reaction *r*_*i*_ and the function *g*_*i*_ defined by (5).
    iv. Calculate the auxiliary values 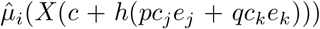 and 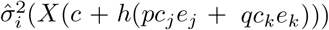 from (7) and, from (6),

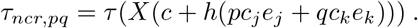
  d. **Find candidate step sizes**, *τ*_*cr,pq*_, **for critical reactions**; for all *p, q* ∈ *{*0, 1*}*:
    i. Calculate 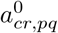 as the sum of the propensities for critical reactions corresponding to the trajectory for *c* + *h*(*pc*_*j*_*e*_*j*_ + *qc*_*k*_*e*_*k*_).
    ii. Choose samples *η*_*pq*_ from the uniform distribution *U*(0, 1) and generate the time to the first critical reaction:

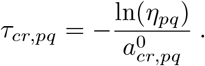
  e. **Calculate the step size**, *τ* : *τ*_*ncr*_ = min_*p,q*∈*{*0,1*}*_*{τ*_*ncr,pq*_ *}, τ*_*cr*_ = min_*p,q*∈*{*0,1*}*_*{τ*_*cr,pq*_ *}, τ* = min(*τ*_*ncr*_, *τ*_*cr*_).
  f. **Calculate the number of reactions** *R*_*ℓ*_, namely *k*_*ℓ*_:
    i. If *R*_*ℓ*_ is a critical reaction, then *k*_*ℓ,pq*_ = 0, for all *p, q* ∈ *{*0, 1*}*.
    ii. If *τ*_*cr*_ ≤ *τ*_*ncr*_, then at least one critical reaction happens. For each *p, q* with *τ*_*cr,pq*_ = *τ*_*cr*_, generate *ℓ*^*^ as a sample of the integet random variable with probabilities 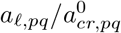, with running only over critical reactions. Put 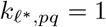.
    iii. For *ℓ* ∈ *J*_*ncr*_, compute *α*_*ℓ,b*_, *b* ∈ *B*, from (12) and sample Poisson random variables *k*_*ℓ,b*_ = Poisson(*α*_*ℓ,b*_*τ*)
    iv. Put: 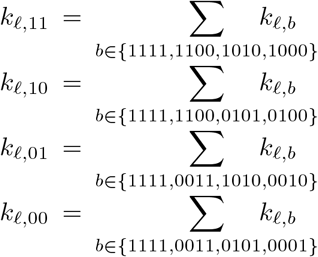
  g. **Update the time and the states on all trajectories**:
    i. *t* = *t* + *τ*.
    ii. 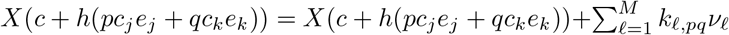, for all *p, q* ∈ *{*0, 1*}*.
  h. **Estimate the second order sensitivity**:

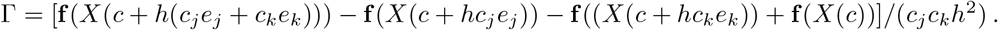

We note that the relative perturbation size *h* is a key parameter, which influences the variance of the sensitivity approximation. In [33, 35], it is shown that the variance of the CFD-2 sensitivity estimator is of order 𝒪(*h*^−3^). In our numerical experiments, we observe a variance of the CTL-2 sensitivity estimator similar to that of CFD-2, both achieving a lower variance, of order 𝒪(*h*^−2^). If the perturbation *h* is too small, the variance of the estimator increases, leading to more noise affecting the sensitivity estimate. On the other hand, if the perturbation *h* is too large, a finite-difference approximation may become inaccurate, while the sensitivity estimates are less noisy. Choosing an appropriate *h* is important for balancing the approximation error of the finite-difference scheme and the variance in the sensitivity estimates, in particular for secondorder ones.

The computational cost of the proposed sensitivity estimation method based on tau-leaping is generally not expected to vary significantly with *h*, when the absolute values of the perturbations are small. The computational cost of estimating these sensitivities depends on the tolerance *ε* of the adaptive tau-leaping scheme. The tolerance *ε* is trade-off parameter which can be used to balance computational efficiency and accuracy of the sensitivity estimation. Significant computational efficiency may be obtained by introducing a small error.

## 4 Numerical Experiments

In this section, we test the efficiency and accuracy of the proposed technique for estimating second-order sensitivities of the Chemical Master Equation, on three models of biochemical networks arising in applications. For each model, we simulate 20,000 Monte Carlo paths with the new method based on tau-leaping, CTL-2, and the existing Coupled-Finite Difference scheme for approximating the Hessian, CFD-2. In each case, we applied a forward finite-difference strategy for estimating sensitivities, with the same relative perturbation *h* of the parameters of interest. We report the evolution in time of the mean trajectories corresponding to the nominal value of the parameter for both methods, and of the standard deviation of these trajectories. We also compare the time-dependence of the estimated second-order sensitivities for both strategies and of the standard deviation of their sensitivity estimators. In addition, we indicate the speed-up of the CTL-2 algorithm over the CFD-2 scheme for each of the three systems. We measure the efficiency gain of our algorithm over the CFD-2 one as

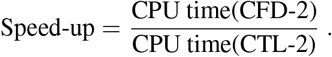

A substantial computational efficiency of the new method compared to the existing scheme is obtained in our tests, which were performed on models with some degree of stiffness.

### 4.1 Decay-Dimerization Model

We start by testing our strategy to estimate sensitivities on the decay-dimerization system of Gillespie [7]. In this model, three molecular species participate in four reactions. The reactions, their propensities and the values of the kinetic parameters are included in Table 1. The system is studied on the time interval [0, 4] with the initial molecular amounts **X**(0) = [10^5^, 10, 0].

**Table 1:** Decay-dimerization model.

|  | Reaction | Propensity | Parameter value |
| --- | --- | --- | --- |
| $R_1$ | $S_1 \xrightarrow{c_1} \emptyset$ | $a_1 = c_1 X_1$ | $c_1 = 1$ |
| $R_2$ | $S_1 + S_1 \xrightarrow{c_2} S_2$ | $a_2 = c_2 X_1(X_1 - 1)/2$ | $c_2 = 0.004$ |
| $R_3$ | $S_2 \xrightarrow{c_3} S_1 + S_1$ | $a_3 = c_3 X_2$ | $c_3 = 0.5$ |
| $R_4$ | $S_2 \xrightarrow{c_4} S_3$ | $a_4 = c_4 X_2$ | $c_4 = 0.04$ |

In our simulations, we apply the CTL-2 method with the value of the tolerance *ε* = 0.05. The mean and standard deviation of the molecular numbers of the species of interest *S*_3_, computed with the CTL-2 and the CFD-2 algorithms are shown in Figures 1(a) and 1(b), respectively. The plots show an excellent match of the means and standard deviations obtained with these strategies, built on the next reaction method (CFD-2) and the adaptive tau-leaping (CTL-2). This shows that the adaptive tau-leaping method utilized by our algorithm is very accurate for the chosen tolerance.

**Figure 1:**
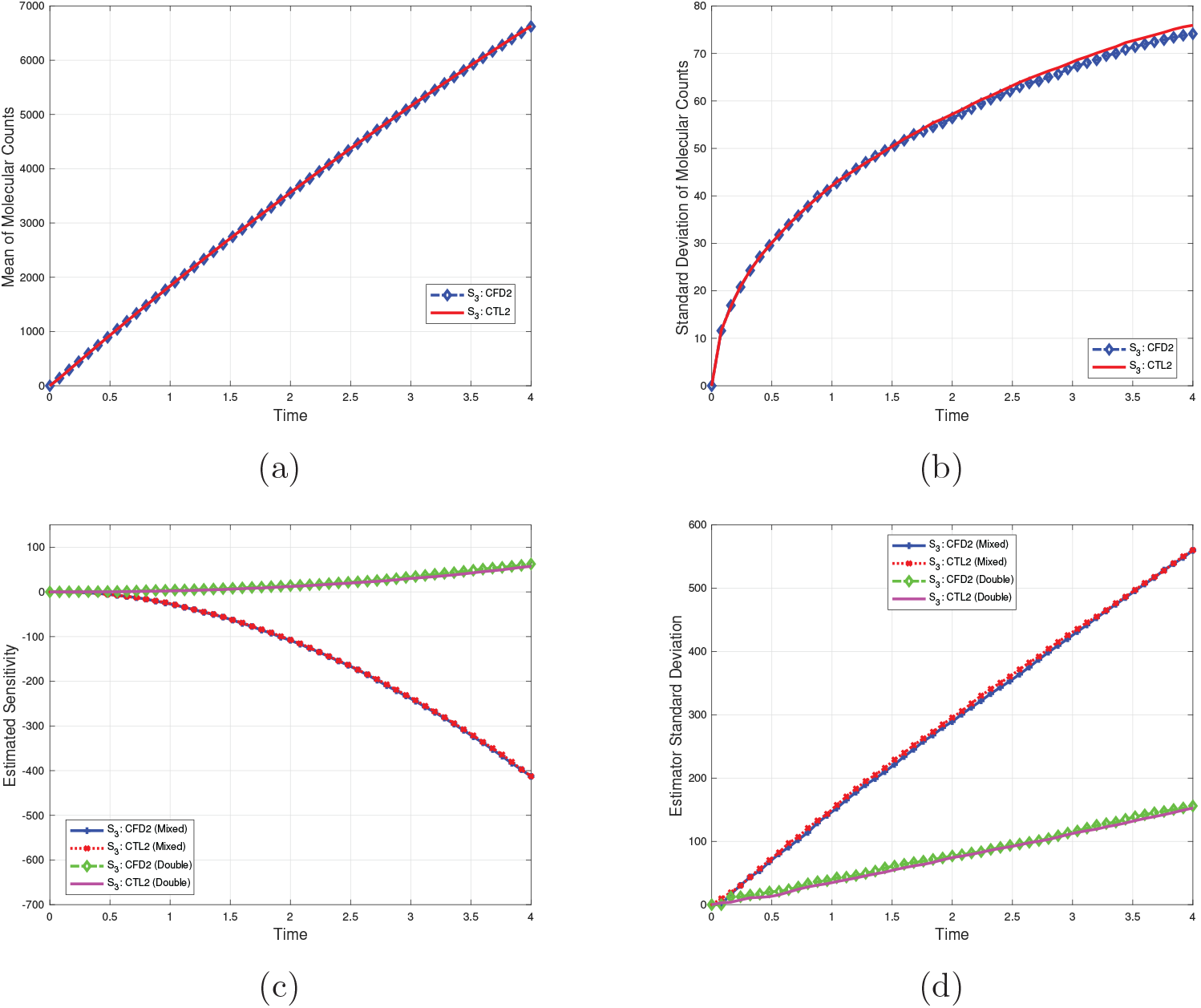
Decay-dimerization model: (a) Mean, (b) Standard deviation of the molecular counts of *S*_3_ by the *τ* - leaping and the NRM methods. (c) Forward finite-difference sensitivity estimators of second-order with respect to *c*_1_ (double) and with respect to *c*_1_ and *c*_3_ (mixed) of the molecular amount of *S*_3_ computed with the CTL-2 and CFD-2 algorithms. (d) Standard deviation of the above CTL-2 and CFD-2 sensitivity estimators. 20,000 quadruples of paths are generated with the CFD-2 and CTL-2 algorithms on the interval [0, 4].

First, let us compute the *double derivative with respect to parameter c*_1_. The relative value of the perturbation is *h* = 0.025, that is 2.5% of the nominal value of the parameter of interest. Figure 1(c) compares the second-order sensitivity with respect to the first parameter as a function of time, 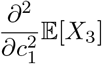 approximated with our method and the CFD-2 scheme. The agreement is excellent, validating our method. In Figure 1(d), a very good match is observed for the evolution in time of the standard deviation of the CTL-2 and the CFD-2 sensitivity estimators, which verifies the accuracy of our technique. Furthermore, we investigate experimentally the dependence of the CFD-2 and CTL-2 sensitivity estimator standard deviations on the perturbation *h*. The findings, provided in Figure 3 (a), show a very good agreement of the standard deviations of the two sensitivity estimators. The plots suggest an order 𝒪 (*h*^−2^) variance of the CFD-2 and CTL-2 methods for this nonlinear model.

Now, let us determine the *mixed derivative with respect to parameters c*_1_ *and c*_3_. In our simulations, the relative value of the perturbation is *h* = 0.05 (i.e., the absolute values of the perturbations are 5% of the parameter values of *c*_3_ and *c*_1_, respectively). The time dependence of the estimated second-order sensitivity of the molecular amount of species *S*_3_ with respect to parameters *c*_1_ and *c*_1_, 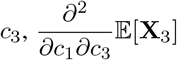, approximated with the CTL-2 and the CFD-2 schemes is given in Figure 1(c). Again, the plots show that our estimator is very accurate. This is confirmed by the excellent agreement obtained for the standard deviations of the CTL-2 and CFD-2 sensitivity estimators, which are plotted versus time in Figure 1(d). When computing the mixed derivative, the computational efficiency of the new method to estimate second-order sensitivities, CTL-2, over the CFD-2 scheme is

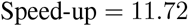

which shows *substantial reductions in computational cost* of our method compared to the existing technique, for a similar accuracy of the sensitivity estimation.

### 4.2 Two-Step Closed Reaction Chain Model

Consider below a two-step closed reaction chain system [36] consisting of three species which interact through four reactions. Table 2 describes the reaction channels and their propensity functions, and specifies the values of the kinetic parameters. This biochemical model is studied with the initial conditions **X**(0) = [2000, 1000, 100], on the time interval [0, 0.1]. The first two reactions are significantly faster compared to the last two ones and, as a consequence, the model is quite stiff.

**Table 2:** Two-step closed reaction model.

|  | Reaction | Propensity | Parameter value |
| --- | --- | --- | --- |
| $R_1$ | $S_1 \xrightarrow{c_1} S_2$ | $a_1 = c_1 X_1$ | $c_1 = 800$ |
| $R_2$ | $S_2 \xrightarrow{c_2} S_1$ | $a_2 = c_2 X_2$ | $c_2 = 3200$ |
| $R_3$ | $S_2 \xrightarrow{c_3} S_3$ | $a_3 = c_3 X_2$ | $c_3 = 0.1$ |
| $R_4$ | $S_3 \xrightarrow{c_4} S_2$ | $a_4 = c_4 X_3$ | $c_4 = 1$ |

We estimate the *double derivative with respect to parameter c*_4_ of the molecular abundance of species *S*_3_, that is 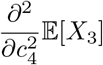. The finite-difference sensitivity estimators are applied with a value of the relative perturbation parameter *h* = 0.05 (the perturbation being 5% of the nominal value of the parameter of interest). We investigated the performance of our tau-leaping second-order sensitivity estimator with the tolerances *ε* = 0.04 and 0.05. Figures 2(a) and 2(b) show the mean and standard deviation of the number of molecules of species *S*_3_ versus time, respectively, computed for the nominal value of the parameter, with the adaptive tau-leaping scheme utilized in the CTL-2 algorithm and the exact modified next reaction method. The agreement of the results produced by these methods is excellent, indicating that the tau-leaping strategy employed by our algorithm is accurate.

**Figure 2:**
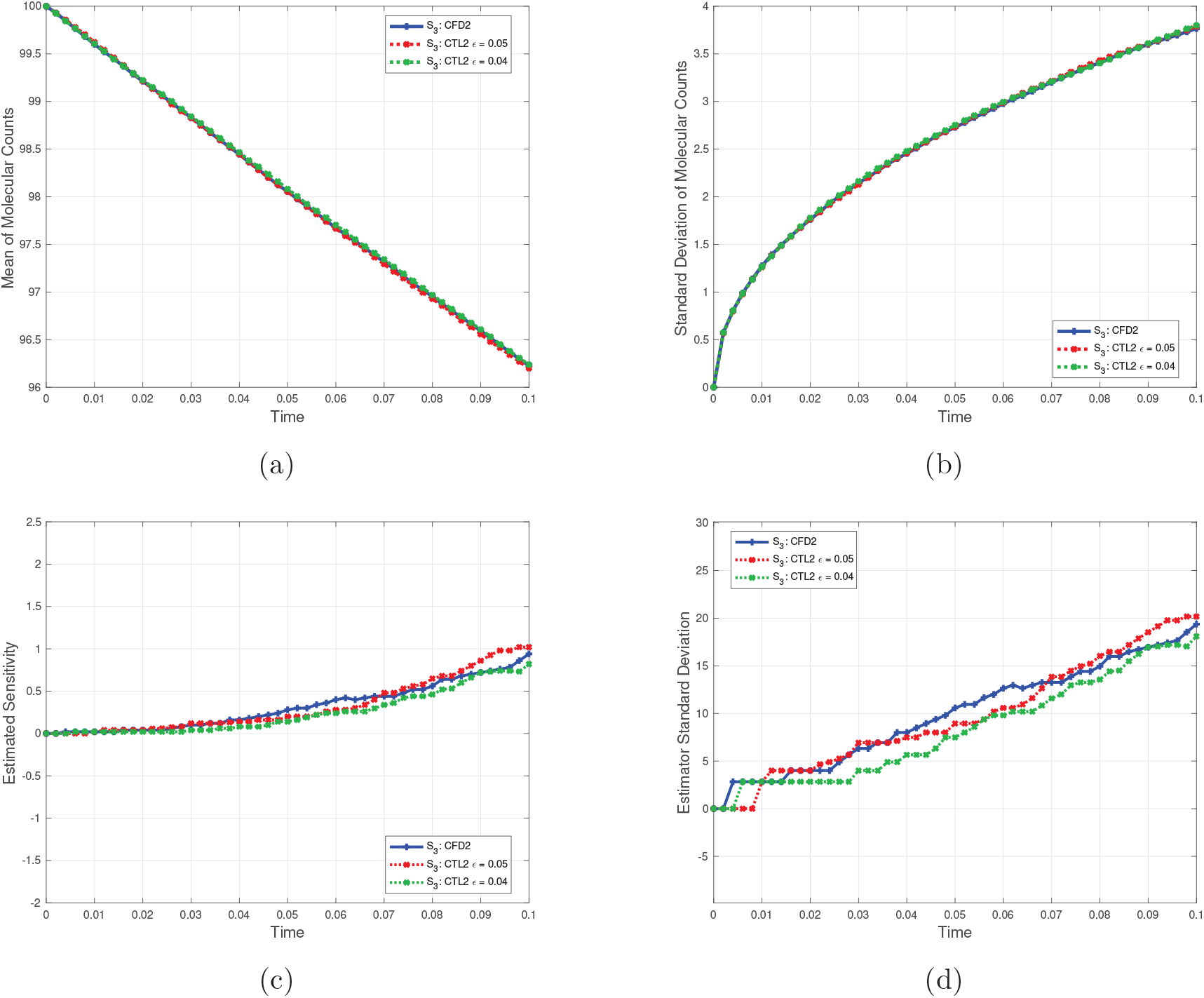
Two-step closed reaction chain model: (a) Mean, (b) Standard deviation of the molecular counts of *S*_3_ by the *τ* -leaping and the NRM methods. (c) Forward finite-difference sensitivity estimators of second-order with respect to *c*_4_ of the molecular amount of *S*_3_ computed with the CTL-2 and CFD-2 algorithms. (d) Standard deviation of the above CTL-2 and CFD-2 sensitivity estimators. 20,000 quadruples of paths are generated with the CFD-2 and CTL-2 algorithms on the interval [0, 0.1].

**Figure 3:**
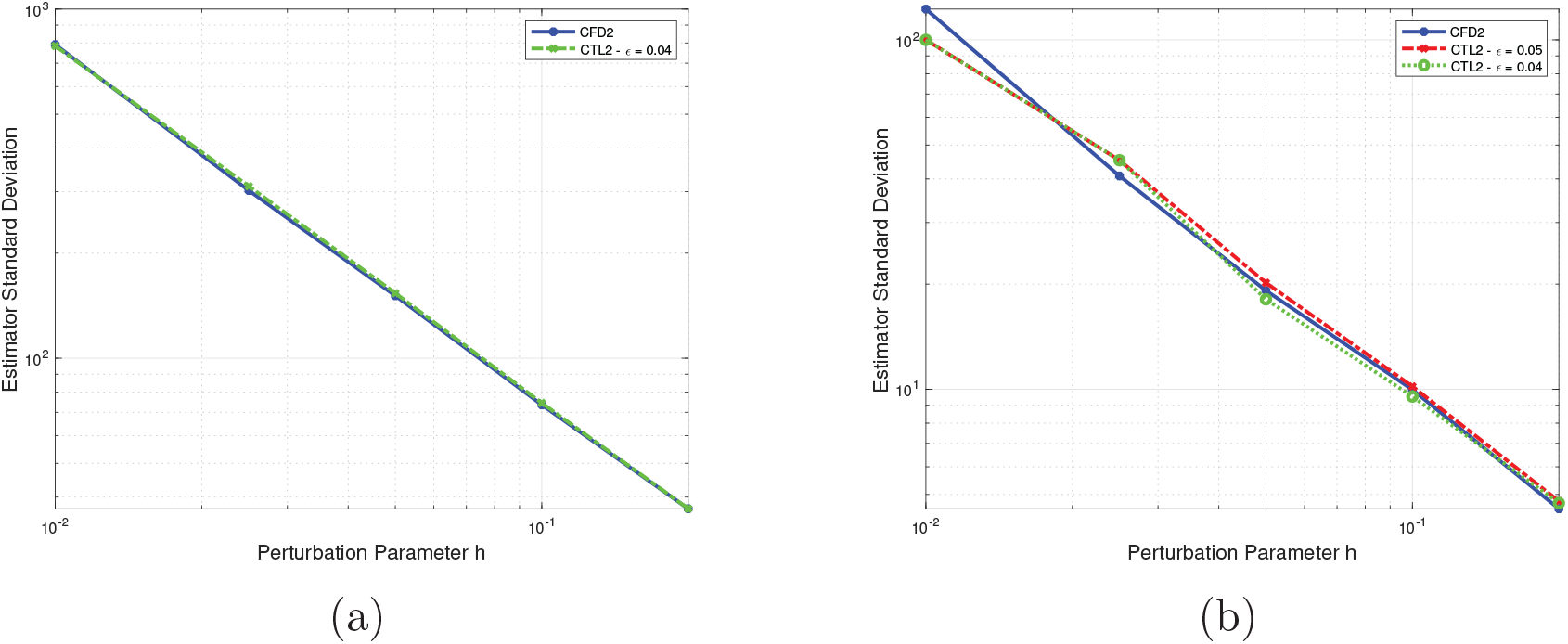
Variation of the standard deviation of the sensitivity estimator of the *S*_3_ molecular counts with respect to *h*, for the CTL-2 and CFD-2 algorithms: (a) the double derivative with respect to *c*_1_ (the decay-dimerization model) and (b) the double derivative with respective to *c*_4_ (the two-step model).

The second-order sensitivity of the number of *S*_3_ molecules with respect to parameter *c*_4_ is estimated by finite-difference schemes with the CTL-2, with tolerances 0.04 and 0.05, and the CFD-2 scheme, their evolution being displayed in Figure 2(c). The results match very well, demonstrating the accuracy of our technique for various tolerances. Furthermore, the standard deviations of the CTL-2 and CFD-2 sensitivity estimators depending on time are in agreement, as shown in Figure 2(d). This further confirms the accuracy of the proposed strategy for estimating sensitivities. Furthermore, the computational efficiency of the CTL-2 algorithm over the CFD-2 is displayed in Table 3. On this model, the new technique has a considerably improved computational efficiency compared to the existing scheme, for a similar accuracy of the sensitivity estimation.

**Table 3:** Two-step closed reaction chain model: speed-up of CTL-2 compared to CFD-2, for estimating the second order sensitivities with perturbation parameter *h* = 0.05. The time interval is [0, 0.01].

| Algorithm | Tolerance | Speed-up |
| --- | --- | --- |
| CTL-2 | $\varepsilon = 0.05$ | 16.09 |
| CTL-2 | $\varepsilon = 0.04$ | 11.72 |
| CFD-2 | – | 1 |

In addition, we analyzed numerically the dependence on the perturbation parameter *h* of the standard deviation of the CFD-2 and CTL-2 sensitivity estimators (for a range of tolerances). The results are displayed in Figure 3 (b). The two algorithms give very similar results, which indicates that the new sensitivity estimator has an accuracy comparable to that of the CFD-2 scheme. Numerically, the variance of the proposed adaptive tau-leaping second-order sensitivity estimator seems to be 𝒪(*h*^−2^) for this model. As mentioned earlier, the smaller values of the perturbation parameter *h* yield more accurate approximations of the true second-order derivative. Yet, these small perturbation sizes increase the variance of the sensitivity estimates, resulting in larger fluctuations in the output. However, large values of the perturbation parameter can introduce considerable truncation errors when approximating the derivative. At the same time, they also lower the variance of the sensitivity estimates, thereby reducing stochastic noise. We determined satisfactory perturbation parameter values experimentally. Exploring the optimal selection of perturbation size, which provides a balance between derivative approximation accuracy and sensitivity estimator variance reduction, falls outside the scope of this paper and remains a subject for future investigation.

The speed-up of our strategy over the existing one, for the tolerance values *ε* tested, is recorded in Table 3. Note that, for a similar accuracy of the sensitivity estimation, the CTL-2 algorithm is *significantly more efficient* compared to the CFD-2 one.

### 4.3 Gene Regulatory Network Model

The third model represents a gene regulatory network [37, 30]. This model was also studied in [38] and, in a different setting not examined here, can display bistable behaviour [39]. For this biochemical network, 8 species interact through 12 reaction channels. Table 4 includes the reactions, their propensities and the values of the reaction rate parameters. The system is analyzed on the time interval [0, 0.01] with initial conditions **X**(0) = [800, 800, 500, 500, 400, 500, 400, 500]^*T*^. This relatively large model is also stiff, with several orders of magnitude separating the fast and slow reactions.

**Table 4:** Gene Regulatory System model.

|  | Reaction | Propensity | Parameter value |
| --- | --- | --- | --- |
| $R_1$ | $S_3 \xrightarrow{c_1} S_3 + S_1$ | $a_1 = c_1 X_3$ | $c_1 = 0.16$ |
| $R_2$ | $S_4 \xrightarrow{c_2} S_4 + S_2$ | $a_2 = c_2 X_4$ | $c_2 = 0.16$ |
| $R_3$ | $S_3 + S_2 \xrightarrow{c_3} S_5$ | $a_3 = c_3 X_3 X_2$ | $c_3 = 5$ |
| $R_4$ | $S_5 \xrightarrow{c_4} S_3 + S_2$ | $a_4 = c_4 X_5$ | $c_4 = 3000$ |
| $R_5$ | $S_5 + S_2 \xrightarrow{c_5} S_6$ | $a_5 = c_5 X_5 X_2$ | $c_5 = 2.5$ |
| $R_6$ | $S_6 \xrightarrow{c_6} S_5 + S_2$ | $a_6 = c_6 X_6$ | $c_6 = 1600$ |
| $R_7$ | $S_1 \xrightarrow{c_7} \emptyset$ | $a_7 = c_7 X_1$ | $c_7 = 0.1$ |
| $R_8$ | $S_4 + S_1 \xrightarrow{c_8} S_7$ | $a_8 = c_8 X_4 X_1$ | $c_8 = 2$ |
| $R_9$ | $S_7 \xrightarrow{c_9} S_4 + S_1$ | $a_9 = c_9 X_7$ | $c_9 = 3000$ |
| $R_{10}$ | $S_7 + S_1 \xrightarrow{c_{10}} S_8$ | $a_{10} = c_{10} X_7 X_1$ | $c_{10} = 2.5$ |
| $R_{11}$ | $S_8 \xrightarrow{c_{11}} S_7 + S_1$ | $a_{11} = c_{11} X_8$ | $c_{11} = 1600$ |
| $R_{12}$ | $S_2 \xrightarrow{c_{12}} \emptyset$ | $a_{12} = c_{12} X_2$ | $c_{12} = 0.1$ |

The numerical simulations are carried out on 20,000 correlated trajectories with the Coupled Finite Difference-2 and the Coupled Tau-Leaping-2 methods, on the time-interval [0, 0.01]. The CTL-2 algorithm is tested with the tolerance values *ε* = 0.05, 0.07 and 0.1. We investigated the accuracy of the tau-leaping strategy on which the CTL-2 scheme is based, by comparing the results with those of the exact Monte Carlo strategy of the Next Reaction Method, for the nominal parameter values. The mean and standard deviations of the molecular numbers of the species *S*_2_ and *S*_5_ as functions of time are displayed in Figures 4(a) and 4(b), respectively. The two strategies lead to similar results and the accuracy of the tau-leaping method is improving with a smaller tolerance *ε*, as expected.

**Figure 4:**
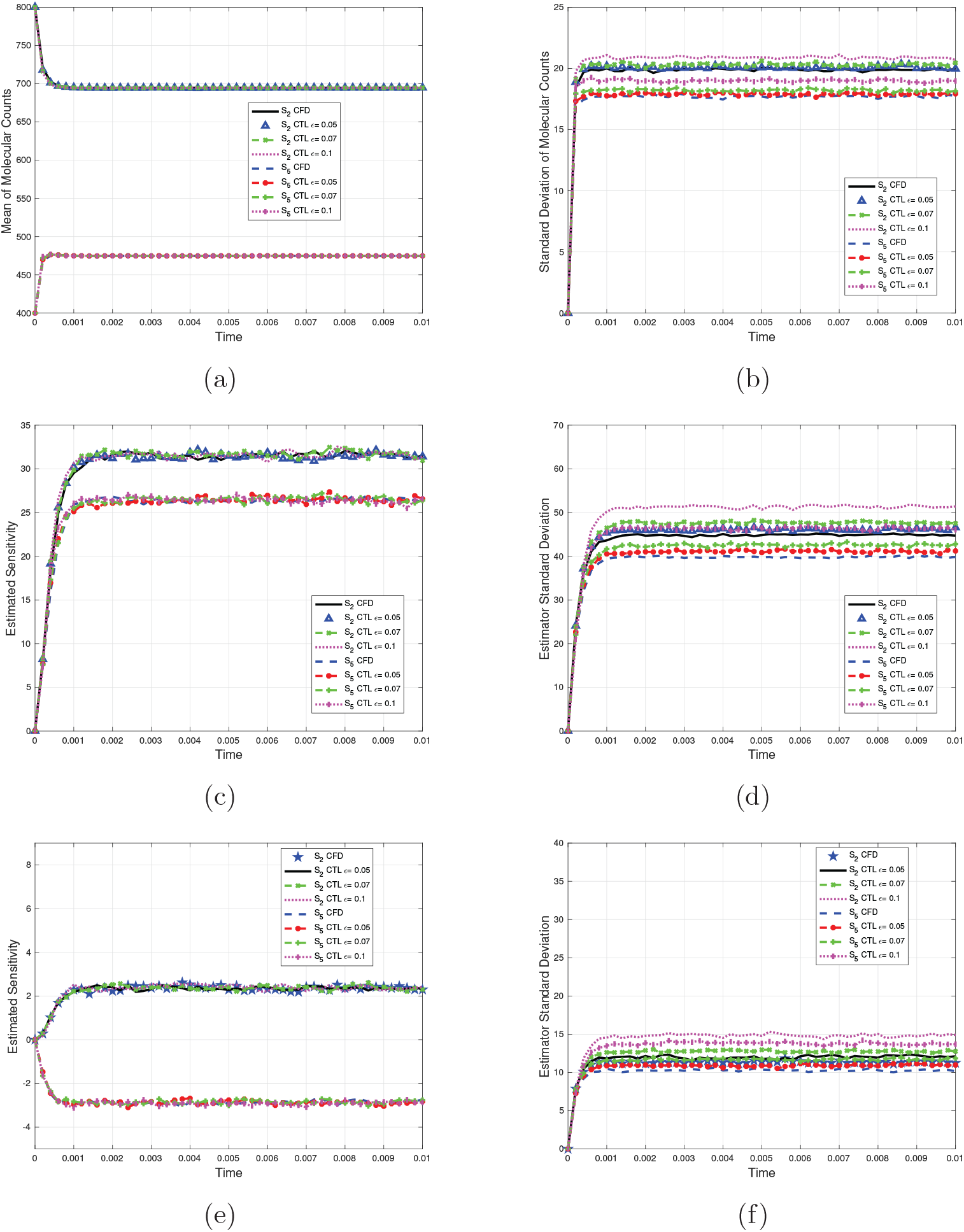
Gene regulatory network model: (a) Mean, (b) Standard deviation of the molecular counts of *S*_2_ and *S*_5_ by the *τ* -leaping and the NRM methods. (c) Forward finite-difference sensitivity estimators of second-order with respect to *c*_5_ and (e) with respect to *c*_3_ and *c*_5_ of the molecular amount of *S*_2_ and *S*_5_, computed with the CTL-2 and CFD-2 algorithms. (d) Standard deviation sensitivity estimators of second-order with respect to *c*_5_ and (f) with respect to *c*_3_ and *c*_5_, of the CTL-2 and CFD-2 algorithms. 20,000 quadruples of paths are generated with the CFD-2 and CTL-2 algorithms on the interval [0, 0.01].

We start by computing the *second-order sensitivity with respect to parameter c*_5_ that is 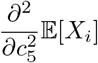 for the indices *i* = 1, …, *N*. The finite-difference sensitivity estimators are employed with the relative perturbation size *h* = 0.05 (a perturbation representing 5% of the parameter value). The time-dependence of the estimated sensitivities of the *S*_2_ and *S*_5_ molecular counts, computed with the CFD-2 and the CTL-2 method with the tolerance values 0.05, 0.07 and 0.1 are shown in Figure 4(c). The agreement is very good for each species. This observation is further verified by comparing the evolution of the standard deviation of the CTL-2 and CFD-2 sensitivity estimators for each of these species, which are plotted in Figure 4(d). These results are similar, with the accuracy of the CTL-2 algorithm depending on the tolerance *ε*.

Finally, we estimate the *second-order sensitivity with respect to parameters c*_3_ *and c*_5_, namely [neq] for each species. The perturbation parameter has the same relative perturbation size as before, *h* = 0.05 (that is 5% of the parameter values of *c*_3_ and *c*_5_, respectively). The second-order sensitivity estimators of the *S*_2_ and *S*_5_ generated with the CTL-2 and the CFD-2 strategies are presented in Figure 4(e), while the standard deviation of their sensitivity estimators are displayed in Figure 4(f). Once more, the similar values of the sensitivity estimators show the accuracy of the CTL-2 method, which is also confirmed in Figure 4(e). Additionally, we measured the computational cost of the two strategies, for a similar level of accuracy. Table 5 displays the speed-up of the tau-leaping sensitivity estimator over the existing one, for several values of the tolerance *ε*. A *significant efficiency gain* of the CTL-2 compared to CFD-2 algorithm is observed for this model of relatively large size, which also exhibits stiffness.

**Table 5:** Gene regulatory network model: speed-up of CTL-2 compared to CFD-2, for estimating the second order sensitivities, with perturbation parameter *h* = 0.8. The time interval is [0, 0.01].

| Algorithm | Tolerance | Speed-up |
| --- | --- | --- |
| CTL-2 | $\varepsilon = 0.05$ | 7.69 |
| CTL-2 | $\varepsilon = 0.07$ | 10.17 |
| CTL-2 | $\varepsilon = 0.1$ | 17.35 |
| CFD-2 | — | 1 |

## 5 Conclusions

Cellular processes frequently involve complex networks of molecular interactions, including enzyme-substrate reactions, molecular binding and dissociation, and biochemical signalling pathways. These systems exhibit stochastic behaviour due to the discrete and probabilistic nature of molecular interactions. Chemical Master Equation is a stochastic discrete model of well-stirred biochemical systems, which accurately captures the randomness and variability observed at the molecular level. Many biochemical systems arising in applications evolve on multiple scales in time, which means that their mathematical models are stiff.

In this study, we proposed a new technique for estimating second derivative sensitivities with respect to various parameters in stochastic discrete models of homogeneous biochemical systems. By accounting for curvature effects and parameter correlations, second-order sensitivity analysis enhances the accuracy and efficiency of parameter estimation, leading to improved model predictions and a better understanding of the underlying biological mechanisms. The proposed finite-difference method to estimate second-order local sensitivities is based on adaptive tau-leaping strategy to compute correlated quartets of trajectories.

The new sensitivity estimator utilizes a strong coupling strategy of the nominal and perturbed trajectories, thereby reducing the estimator variance. The tau-leaping framework for second-order sensitivities combines variance reduction with computational efficiency. This approach is valuable not only for its effectiveness, but also for making second-order sensitivity analysis feasible for large and complex biochemical systems, where exact methods would be computationally prohibitive.

We successfully tested our strategy on several models of biochemical networks, from small to relatively large size systems. The proposed Coupled Tau-Leaping-2 technique, is significantly more efficient than the existing Coupled Finite Difference-2 scheme, for a similar accuracy, on biochemical models which are moderately stiff to stiff. In the future, we plan to study the dependence of the accuracy of the sensitivity estimator on the perturbation parameter.

## Acknowledgements

This research was supported by a grant from the National Sciences and Engineering Research Council of Canada (NSERC) - Grant No. RGPIN-2020-05469, and Toronto Metropolitan University.

## Notes

### Competing Interest Statement

The authors have declared no competing interest.

